# Direct and abscopal anti-tumor responses elicited by peripheral nerve schwannoma irradiation synergize with anti-PD1 treatment in vestibular schwannoma models

**DOI:** 10.1101/2024.12.29.630655

**Authors:** Zhenzhen Yin, Simeng Lu, Limeng Wu, Yao Sun, David W. Blake, Jie Chen, Lukas D. Landegger, William Ho, Bingyu Xiu, Adam P. Jones, Alona Muzikansky, Helen A. Shih, Konstantina M. Stankovic, Scott R. Plotkin, Lei Xu

## Abstract

*NF2*-related schwannomatosis (*NF2*-SWN) is a progressive and disabling disease requiring effective treatments. The hallmark of *NF2*-SWN is bilateral vestibular schwannomas (VSs), which progressively enlarge, leading to permanent sensorineural hearing loss and severely impacting patients’ quality of life. Currently, there are no FDA-approved drugs for VS or the associated hearing loss. Immune checkpoint inhibitors (ICIs) have revolutionized cancer treatment, but have not yet been systematically investigated in non-malignant tumors such as VS. In our studies, we demonstrated that combining anti-PD1 (αPD1) treatment with radiation therapy (RT) provides three significant therapeutic benefits: i) Enhanced αPD1 efficacy and immune memory: RT induces immunogenic cell death and activates the STING pathway, enhancing αPD1 efficacy and generating long-term immune memory, ii) Reduced RT dose and associated tissue injury: The combination strategy reduces the required RT dose necessary for effective tumor control, potentially minimizing RT injury to surrounding normal tissues, and iii) Elicited abscopal effects on cerebellopontine angle (CPA) schwannomas: RT to peripheral nerve tumor induces a systemic abscopal effect, which synergizes with αPD-1 to effectively control intracranial schwannomas without direct irradiation, sparing the cochlea from radiation exposure and avoiding auditory radiation injury. Together, our findings provide a compelling rationale for deploying ICIs in combination with radiotherapy as a novel treatment approach for patients with VS and *NF2*-SWN.

## Introduction

*NF2*-related schwannomatosis (*NF2*-SWN) is a dominantly inherited neoplasia syndrome caused by germline pathogenic variants in the *NF2* tumor suppressor gene. *NF2*-SWN has an incidence of 1 in 61,000 people and a penetrance of nearly 100% ^[1, 2]^. *NF2*-SWN patients develop multiple peripheral nerve sheath tumors, and the pathognomonic hallmark is non-malignant, bilateral VSs arising along cranial nerve VIII. Progressively growing VSs lead to sensorineural hearing loss (SNHL) and substantial morbidity, and severely impacting patients’ quality of life ^[3]^. Large VSs can compress the brainstem resulting in severe morbidity and even death ^[4]^. Standard treatments for growing VSs include surgery and radiotherapy (RT), however, both carry risks of damaging the nerve and causing profound deafness, chronic dizziness, and facial and other cranial nerve palsies ^[5–7]^. Identifying effective drugs to halt VS growth and prevent VS-associated SNHL is a major unmet clinical need.

In patients with sporadic VS, tumor control after irradiation is excellent, with long-term tumor control rates of around 88-91%, but significantly lower for patients with *NF2*-SWN ^[8, 9]^. In addition, radiotherapy can cause adverse effects, including: i) further hearing loss due to ototoxicity ^[8, 9]^, ii) pseudoprogression - a transient increase in tumor volume that occurs in 23-44% of VS patients and can worsen brainstem compression ^[10–12]^, and iii) increased risk of malignant transformation ^[13–15]^. There is a need for a combination regimen that can lower the therapeutic dose of RT required for tumor control, and, consequently, reduce radiation-induced ototoxicity and adverse effects.

Immune checkpoint inhibitors (ICIs), such as monoclonal antibodies against programmed cell death protein-1 (PD-1), PD ligand 1 (PD-L1), and cytotoxic T-lymphocyte-associated protein 4 (CTLA-4), block immune inhibitory signals to augment T cell activation and effector functions, leading to enhanced antitumoral immunity and improved prognostic outcomes ^[16, 17]^. ICIs are FDA-approved for a wide range of solid tumors, as monotherapy or in combination with other agents like chemotherapy ^[18]^. However, ICI’s application in non-malignant schwannomas and their therapeutic potential on hearing preservation remain unexplored.

VS harbor substantial numbers of T cells ^[19]^, however, a large fraction of these tumor-infiltrating CD4^+^ and CD8^+^ T cells express PD-1 ^[20]^, and present with a transcriptome signature associated with CD8^+^ T cell senescence ^[21]^. There is high expression of PD-1 and its ligand B7-H1 in VS ^[19, 22]^. Despite these observations, few studies have explored the treatment potential of ICIs in VS *-* anti-PD1 (αPD1) antibody modestly delayed schwannoma growth in a subcutaneous mouse model ^[23]^ and αPD1 salvage therapy led to tumor growth arrest in one patient with recurrent VS ^[24]^. This information suggests that ICIs may be promising treatments for VS.

In this study, we aim to address three questions: 1) can the combination of αPD1 and RT treatment enhance anti-tumor efficacy compared to each monotherapy? 2) can the addition of αPD1 treatment reduce the required RT dose for tumor control, thereby minimizing potential RT-related ototoxicity and other adverse effects? And 3) can local irradiation of a peripheral nerve schwannoma elicit an abscopal effect that synergizes with systemic αPD1 treatment and shrinks intracranial vestibular schwannoma without direct irradiation, thus sparing the cochlea from RT-related ototoxicity and better preserving hearing function?

## Materials and Methods

### Cell lines

Mouse *Nf2*^-/-^ Schwannoma cells were maintained in 10% fetal bovine serum (FBS)-containing Schwann cell medium with Schwann cell growth supplement (SCGS, ScienCell) ^[25]^. Mouse SC4 Schwannoma cells (gift from Dr. Vijaya Ramesh, Massachusetts General Hospital, MGH) were maintained in 10% FBS-containing Dulbecco’s Modified Eagle Medium (DMEM, Corning) ^[26, 27]^.

### Animal models

All animal procedures were performed following the Public Health Service Policy on Humane Care of Laboratory Animals and approved by the Institutional Animal Care and Use Committee of the MGH. *Nf2^-/-^* and SC4 tumors were implanted in immune-competent C57/FVB mice. In all animal experiments, to achieve appropriate statistical power and detect any sex-related differences, we used 8-12 weeks-old mice. Since VS developed in both sexes, we used mice in an equal ratio of male and female mice (1:1) and used age and sex-matched mice under each experimental condition. Because schwannomas grow in the vestibular nerve and peripheral nerves in patients, we used two mouse models by injecting tumor cells into the cerebellopontine angle (CPA) and in the sciatic nerve of mice.

#### Cerebellopontine angle (CPA) model

To recapitulate the intracranial microenvironment of VSs, tumor cells were implanted into the CPA region of the right hemisphere ^[28, 29]^. A total of 1 μl of tumor cell suspension (2,500 cells) was implanted per mouse.

#### Sciatic nerve schwannoma model

To reproduce the microenvironment of peripheral schwannomas, we implanted tumor cells into the mouse sciatic nerve ^[26]^. A total of 3 μl of tumor cell suspension (5x10^4^ cells) was injected slowly (over 45-60 seconds) under the sciatic nerve sheath using Hamilton syringe to prevent leakage.

### Treatment protocols

In the sciatic nerve model, treatment starts when the tumor reaches 3 mm in diameter. In the CPA model, treatment starts when the blood Gaussia luciferase reporter gene (Gluc) reaches 1x10^4^ RLU.

#### Anti-PD1 treatment

The anti-PD1 antibody or isotype control IgG (200 μg/mice, Bioxell) was administrated *i.p.* every 3 days for a total of 4 dosages.

#### Radiation therapy

Treatment was delivered in a ^137^Cs gamma irradiator for small animals, which produces 1.176 MeV gamma rays and allows longitudinal radiation studies. Mice were irradiated locally in a custom-designed irradiation chamber, which shields the entire animal except for the brain (in the CPA model) or the leg (in the sciatic nerve model) ^[26]^. We used 5 Gy in our study as this produced tumor growth delay with the least amount of toxicity (evaluated by body weight loss)^[26]^.

### Measurement of tumor growth

To monitor intracranial CPA tumor growth, both tumor cell lines were infected with lentivirus encoding secreted Gluc reporter gene, and plasma Gluc was measured as previously described ^[29–32]^. Briefly, 13 μl of whole blood was collected from a slight nick on tail vein and mixed with 5 μl 50 mM EDTA immediately to avoid clotting. Blood samples were transferred to a 96-well plate and Gluc activity was measured using a plate luminometer (GloMax 96 Microplate Luminometer, Promega). The luminometer was set to automatically inject 100 µl of 100 mM coelenterazine (CTZ, Nanolight) in phosphate buffered saline (PBS), and photon counts were acquired for 10 sec. Sciatic nerve tumor size was measured using a caliper every 3 days until tumors reached 1 cm in diameter.

### Audiometric testing in animals

Distortion product otoacoustic emissions (DPOAEs) and auditory brainstem responses (ABRs) were measured as described previously ^[29]^. Briefly, animals were anesthetized via i.p. injection of ketamine (0.1 mg/g) and xylazine (0.02 mg/g). The tympanic membrane and the middle ear were microscopically examined for signs of otitis media. All animals had well-aerated middle ears. DPOAEs were recorded in response to two primary tones (f1 and f2) with the frequency ratio f2/f1 = 1.2 at half-octave steps from f2 = 5.66–45.25 kHz while increasing intensity in 5 dB steps from 15 to 80 dB sound pressure level (SPL). The 2f1-f2 DPOAE amplitude and surrounding noise floor were monitored, and the threshold was defined as the f2 intensity that created a distortion tone > 0 dB SPL.

ABRs were recorded between subdermal needle electrodes: positive in the inferior aspect of the ipsilateral pinna, negative at the vertex, and ground at the proximal tail. The responses were amplified (10,000X), filtered (0.3–3.0 kHz), and averaged (512 repetitions) for each of the same frequencies and sound levels as used for DPOAE measurements. Custom LabVIEW software for data acquisition was run on a PXI chassis (National Instruments Corp). For each frequency, the auditory threshold was defined as the lowest stimulus at which repeatable peaks could be observed on visual inspection. In the absence of an auditory threshold, a value of 85 dB was assigned (5 dB above the maximal tested level).

### Analysis of tumor-infiltrating immune cells

To characterize the tumor-infiltrating immune cells, tumor lysates were washed with PBS and stained for flow cytometry with anti-CD45 (clone 30-F11), anti-CD4 (clone RM4-5), anti-CD8 (clone 53-6.7), anti-Foxp3 (clone D6O8R), anti-NK1.1 (clone PK136), anti-Gr1 (clone RB6-8C5), anti-CD11b (clone m1/70), anti-F4/80 (clone BM8), anti-iNOS (clone CXNFT), anti-Arginase I (clone D4E3M), anti-CD86 (clone GL-1), anti-interferon (IFN-γ, clone XMG1.2), anti-TNFα (clone MP6-XT22), and anti-Granzyme B (clone GB11), see Supplementary Table 1 for antibody details.

### Gene expression analysis

#### RNASeq

Fresh mouse tumor samples were homogenized using the polytron PT1300 tissue homogenizer, followed by additional homogenization using a Qiashredder spin column. RNA from tumor tissues was extracted using the RNeasy Mini Kit (QIAGEN, Cambridge, MA). 1 μg of total RNA was sent to Molecular Biology Core Facilities, Dana-Farber Cancer Institute. RNASeq analysis was performed following the routine procedure ^[27]^. The DESeq2 package in R was used to determine the differentially expressed genes (DEGs) ^[33]^. To control for the False discovery rate (FDR) at 0.05, we used the Benjamini & Hochberg correction. ComplexHeatmap package was used to plot the heatmap ^[34]^. The differentially expressed gene set was analyzed by Gene Set Enrichment Analysis software (GSEA, https://software.broadinstitute.org/software/cprg/?q=node/14).

#### Quantitative RT-PCR

qPCR was performed to evaluate changes in mRNA level using SYBR Green-based protocol ^[35]^. All qPCR and analysis were performed on a Stratagene MX 3000 qPCR System operating MXPro qPCR software (Stratagene) ^[36]^.

#### Western Blot

Thirty micrograms of protein per sample were separated on 10% SDS-polyacrylamide gels ^[37]^. Membranes were blotted with antibodies against total (1:500) and phospho-STING (1:1000); total (1:500) and phospho-TBK1 (Ser172, 1:1,000); total (1:500) and phospho-IRF3 (Ser396, 1:1,000); HMGB1 (1:500) and cGAS (1:500). Antibodies were obtained from Cell Signaling (Danvers, MA). Membranes were blotted with antibodies against beta-actin for equal loading control (1:5,000, Sigma) ^[38]^.

#### ELISA

Plasma or protein extracted from snap-frozen tumors were diluted to 2 μg/μl concentration according to protein assay. Mouse inflammatory cytokine levels were quantified using mouse multiplex enzyme-linked immunosorbent assay plates following the manufacturer’s instructions (Meso-Scale Discovery, Gaithersburg, MD). Every sample was run in triplicate ^[35, 39]^.

### *In vitro* viability and apoptosis assay

Cell viability was determined *in vitro* by MTT assay ^[40]^. In vitro apoptosis was determined by Annexin V-FITC and propidium iodide (PI) staining followed by flow cytometry, following the manufacturer’s instruction (Biolegend).

### ATP release assay

ATP release was evaluated using Luminescent ATP Detection Kit (Abcam) following the manufacture’s instruction. Briefly, 50 µL of detergent was added to lyse the cells and stabilize the ATP. The plate was sealed and shaken for 5 minutes at 600-700 rpm. Then, 50 µL of substrate solution was added to each well, and luminescence was measured using a microplate reader. ATP levels were calculated by comparing luminescence values to a standard curve prepared using known concentrations of ATP.

### Histological staining

To evaluate tumor cell proliferation and apoptosis, slides of tumor tissues were stained with proliferating cell nuclear antigen (PCNA, 1:1,000, Abcam) and TUNEL (ApopTag®, EMD Millipore). Archived paraffin-embedded patient VS samples were stained with antibodies against phosphorylated TBK (1:10, Cell Signaling Technology). Four specimens of normal peripheral nerve were obtained postmortem and used as controls. Appropriate positive and negative controls were used for all stains ^[39]^. Histological analysis using digital quantitative image analysis was performed using the open-source software ImageJ. Positive staining in 20 random fields/slides was quantified via automated built-in functions based on fluorescent pixel intensity after establishing a threshold to exclude background staining.

### Statistical analyses

We determined whether growth curves significantly differed from each other by log-transforming the data, fitting a linear regression to each growth curve, and comparing the slopes of the regression lines (using F-test from ANCOVA). Survival analyses and curves were done using the KM method. Differences in tumor growth between the two groups were analyzed using the Student’s t-test (two-tailed) or Mann-Whitney *U* test (two-tailed). For gene expression comparison, significant differences were determined by mixed-effect models to take into account the correlation structure. P values were multiple-test corrected using the Benjamini-Hochberg False Discovery Rate (FDR) procedure. Benjamini-Hochberg correction for multiple comparisons was applied. All calculations were done using GraphPad Prism Software 6.0 and Microsoft Excel Software 2010.

## Results

### Combined radiotherapy and αPD1 show improved tumor control efficacy in both schwannoma models

To address our first question of whether combined RT+αPD1 treatment can achieve enhanced efficacy, we treated mice bearing *Nf2^-/-^* tumors in both sciatic nerve and CPA models with: i) control IgG, ii) αPD1, iii) 5Gy RT, or iv) αPD1+5Gy RT (Figure 1A). In the sciatic nerve model, both RT and αPD1 monotherapy modestly delayed tumor growth, and combined RT+αPD1 treatment was significantly more effective in tumor growth inhibition compared to each monotherapy (Figure 1B). In the CPA model, combined RT+αPD1 treatment significantly prolonged survival - with 62.5% of mice surviving ≥100 days, compared to 32% of long-term survivor mice in the αPD1 monotherapy group (Figure 1C). This enhanced efficacy from the combination treatment was also observed in a second schwannoma, SC4 model (Supplemental Figure 1A).

**Figure 1.**
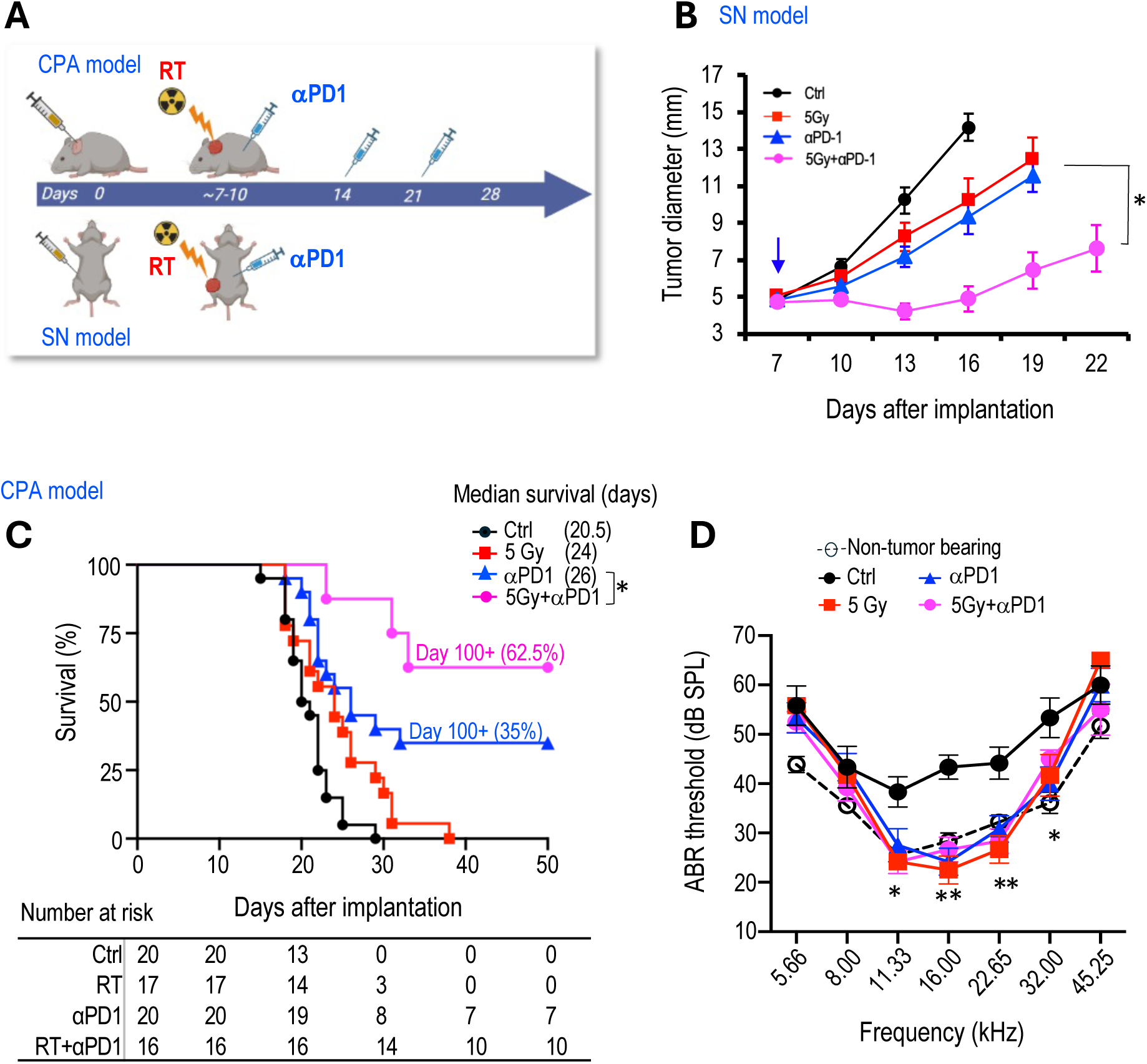
Combined radiotherapy and αPD1 show improved tumor efficacy in both schwannoma models. **(A)** Schematic and timeline of combined radiation therapy and αPD1 treatment in the CPA and SN schwannoma models. **(B)** Tumor diameter measured by caliper in the SN schwannoma model. Arrow, treatment start time. **(C)** Kaplan-Meir survival curve of mice bearing *Nf2^-/-^* tumor in the CPA model. **(D)** ABR threshold of mice bearing *Nf2^-/-^* tumor in the CPA model 21 days post-treatment. All animal studies are presented as mean±SEM, N=6 mice/group, and representative of at least three independent experiments. In vivo study significance was analyzed using a Student’s t-test. *P<0.01, **P<0.001.

The effects of immunotherapy on hearing function remain to be systemically evaluated. We first evaluated potential ototoxicity from αPD1 treatment and from 5 Gy RT to the CPA in non-tumor bearing mice by measuring auditory brainstem evoked response (ABR), which represents the summed activity of the auditory nerve and central auditory nuclei. In non-tumor bearing mice, neither αPD1, 5 Gy RT, nor their combination changed the ABR threshold in mice 42 days post-treatment (Supplemental Figure 1B-D). In mice bearing *Nf2^-/-^* tumors in the CPA region, treatment with αPD1, 5 Gy RT, or the combination of both restored the ABR threshold to normal levels observed in non-tumor-bearing mice. No treatment demonstrated superiority over the others (Figure 1D).

### Combined radiotherapy and αPD1 treatment generate immunologic memory

To test if long-term survivors develop durable immune memory, we re-challenged ‘cured’ mice (the initial tumor completely disappeared as confirmed by negative Gluc measurement, and mice survived over 100 days) with injections of *Nf2^-/-^*-Gluc cells into the CPA region contralateral to the previous injection (Figure 2A). Naïve mice were included as controls for tumor take. In naïve mice, 100% of mice grew tumors, and their blood Gluc reporter gene levels reached over 1x10^6^ RLU by day 14 after implantation. On the contrary, none of the ‘cured’ mice developed tumors by day 63 after the rechallenge (Figure 2B). These data suggest that the ‘cured’ mice have long-term tumor-specific immunological memory against the *Nf2^-/-^* cells.

**Figure 2.**
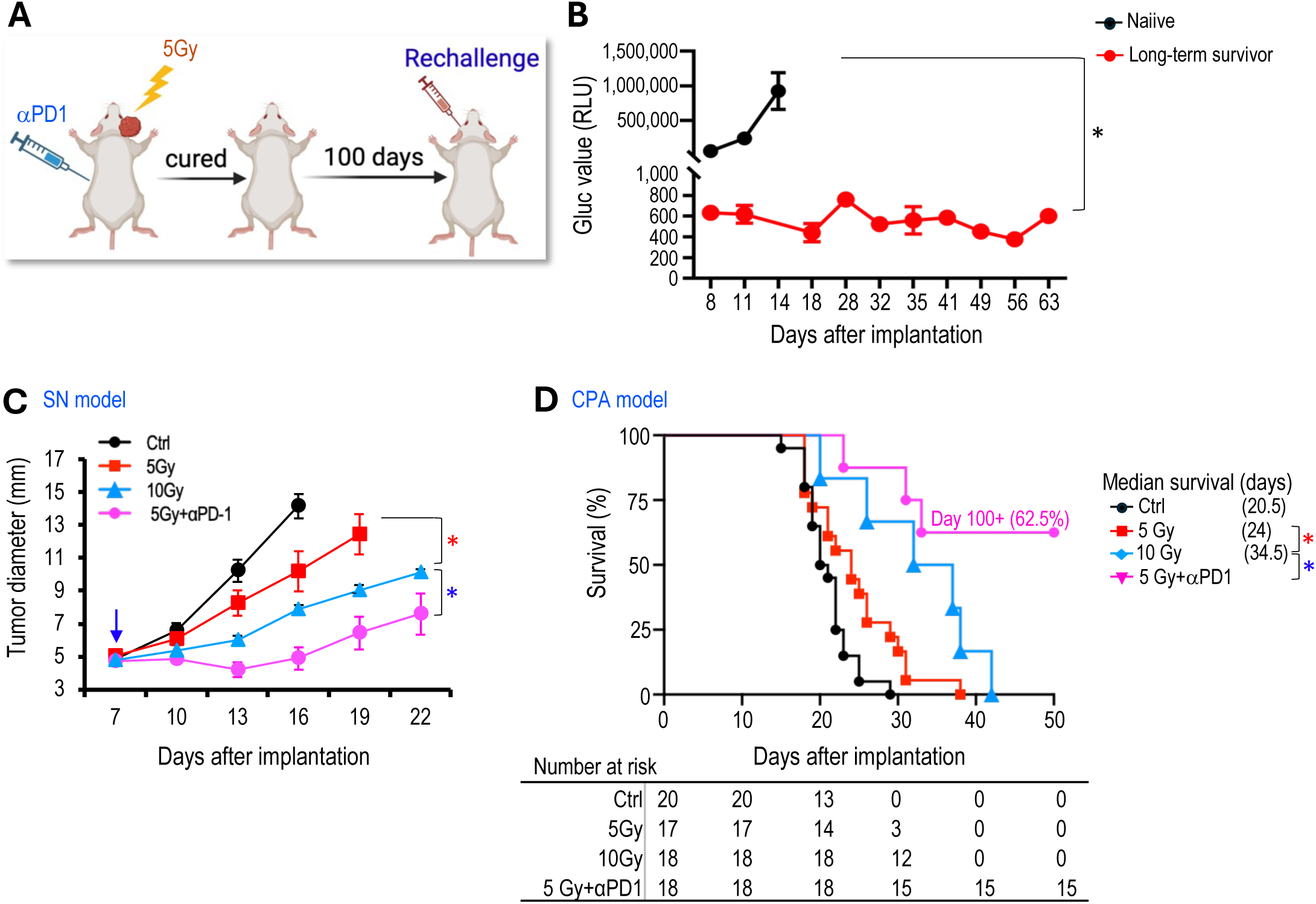
Combined radiotherapy and αPD1 treatment generate immunologic memory and lowered the radiotherapy dose required for tumor control. **(A)** Schematic and timeline of rechallenging experiment in long-term survivors. **(B)** Blood Gluc level in naïve and long-term survivors. **(C)** Tumor diameter measured by caliper in the SN schwannoma model. **(D)** Kaplan-Meir survival curve of mice bearing *Nf2^-/-^* tumor in the CPA model. All animal studies are presented as mean±SEM, N=8 mice/group, and representative of at least three independent experiments. In vivo study significance was analyzed using a Student’s t-test. *P<0.01.

### Combined αPD1 treatment lowered the radiotherapy dose required for tumor control

Key determinants of hearing loss after RT are the dose of radiation and volume of cochlea irradiated, with less radiation associated with better hearing preservation ^[41]^. To address our second question, whether combined αPD1 treatment helps lower the required RT dose for tumor control, thereby minimizing potential RT-related ototoxicity and adverse events, we treated groups of mice with i) control IgG, ii) 5 Gy RT, iii) 10 Gy RT, or iv) 5Gy RT+αPD1. In both sciatic nerve (Figure 2C) and CPA (Figure 2D) models, we found that high-dose RT (10 Gy) was more effective than low-dose RT (5 Gy) in delaying tumor growth and prolonging survival. However, when combined with αPD1 treatment, 5 Gy RT was more effective compared to 10 Gy RT alone. These data suggest that combining αPD1 treatment with RT could help lower the required RT dose for tumor control.

### RT induces immunogenic cell death in schwannoma models

Using RNASeq analysis, we found that irradiation of *Nf2^-/-^*tumor led to a significant upregulation of genes involved in the pyroptosis process (Figure 3A, 3B). Pyroptosis is a form of immunogenic cell death (ICD) sufficient to trigger a systematic immune response ^[42]^,^[43]^. Irradiation of *Nf2^-/-^* cells *in vitro* caused a significant increase in tumor cell pyroptosis and necrosis (Figure 3C, 3D). RT induced the extracellular release of pyroptosis markers: i) high-mobility group box 1 (HMGB1), which is a ‘danger’ signal (Figure 3E), and ii) adenosine triphosphate (ATP), which is a ‘find me’ signal (Figure 3E). Both HMGB1 and ATP are released under conditions that cause pyroptosis ^[44, 45]^. RT also caused tumor cells to express significantly elevated levels of inflammatory cytokines, including, IFN-α, TNF-α, IL-1β, IL-6, and IFN-γ (Figure 3G).

**Figure 3.**
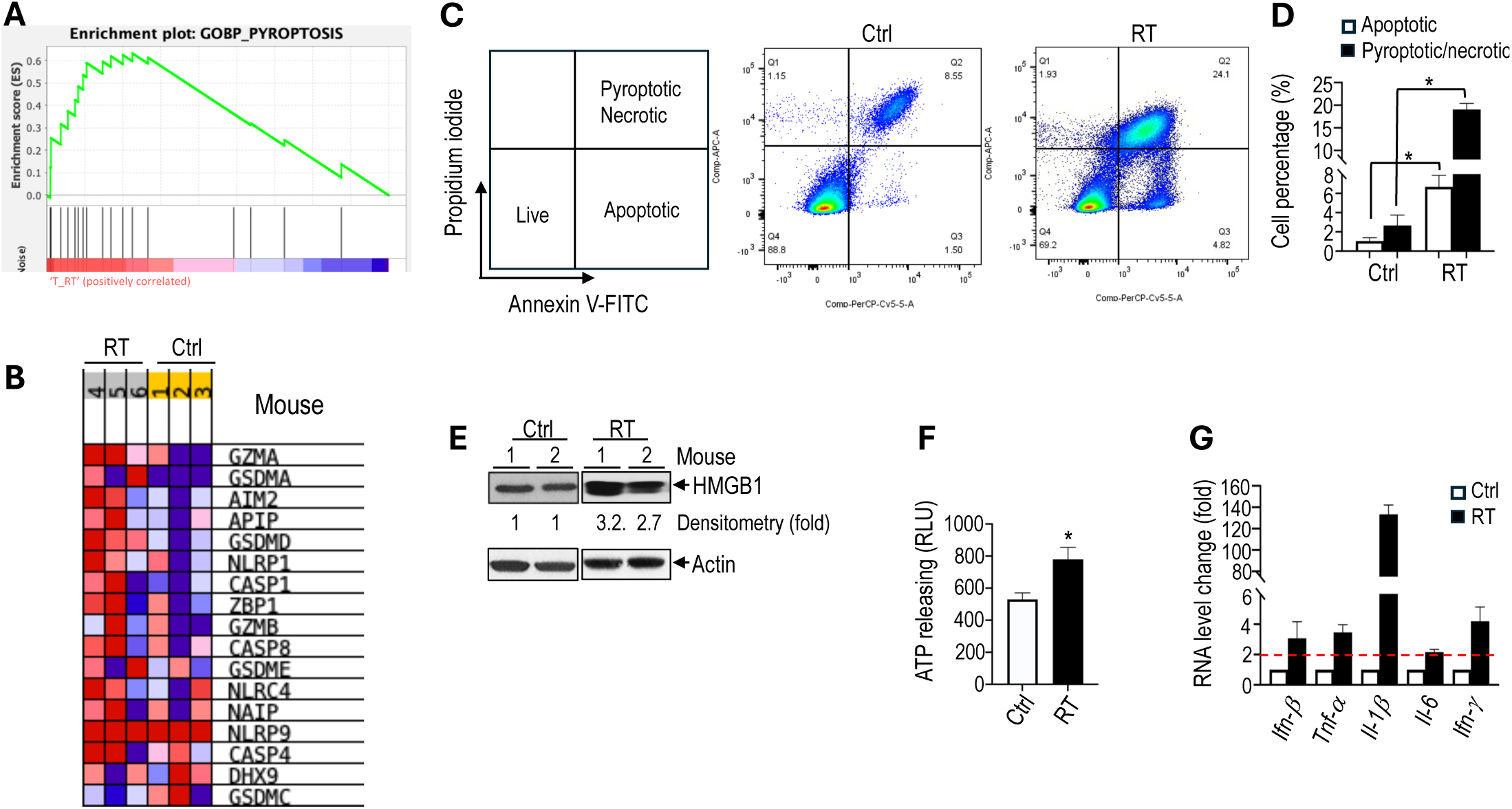
RT induces immunogenic cell death in schwannoma models. RNAseq analysis of bulk tumors from control and irradiated *Nf2^-/-^*tumors from the CPA mouse model (N=3 tumors/group). **(A)** GSEA enrichment plots of GO_Pyroptosis pathway related genes. **(B)** Heatmap of pyroptosis pathway genes. Flow cytometry of Annexin V/Propidium iodide-stained control and 5 Gy in vitro irradiated *Nf2^-/-^* cells. **(C)** Flow cytometry gating of Annexin V/Propidium iodide. **(D)** Flow cytometry analysis of apoptotic or pyroptotic and necrotic *Nf2^-/-^* cells. Pyroptosis-related gene expression analysis. **(E)** Western blot of control and 5 Gy irradiated *Nf2^-/-^* tumor tissues (N=2 tumors/group). **(F)** Fluorescent intensity of extracellular ATP release from control and in vitro 5 Gy irradiated *Nf2^-/-^* cells (n=6 plates/group). **(G)** qRT-PCR analysis of murine *Ifn-β, Tnf-α, Il-1β, Il-6 and Ifn-γ* mRNAs in control and 5 Gy irradiated *Nf2^-/-^* cells (N=3 plate of cells/group). Flow cytometry and gene expression data are presented as mean±SD, and analyzed using Student’s t-test and the Mann-Whitney test. *P<0.01.

### RT of peripheral nerve schwannoma elicits an abscopal effect and synergizes with αPD1 treatment

We next performed an experiment to address our third question: whether local irradiation of a peripheral nerve schwannoma induces an abscopal effect that synergizes with systemic αPD1 treatment, leading to the shrinkage of intracranial vestibular schwannoma without direct irradiation. This approach aims to spare the cochlea from RT-related ototoxicity and better preserve hearing function. We implanted mice with *Nf2^-/-^* cells bilaterally in the sciatic nerves. When tumors grow to ∼5 mm in diameter, mice were randomized into groups receiving control IgG or treatment with i) αPD1 only, or ii) αPD1+5 Gy RT applied only to the tumor on the right leg, while the rest of the body was shielded (Figure 4A). Results indicated that while the group with the tumor receiving both direct RT and αPD1 had the best response (green line), the contralateral non-irradiated tumor receiving systemic αPD1 treatment plus the RT abscopal effect (blue line) showed significantly reduced growth compared to tumors receiving αPD1 alone (red line, Figure 4B).

**Figure 4.**
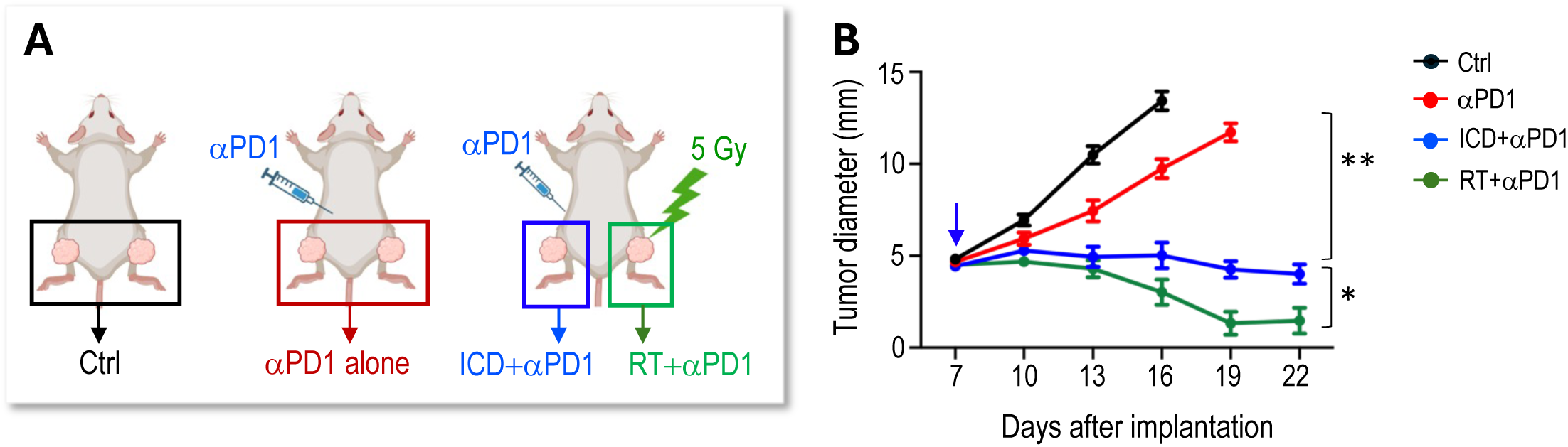
RT of peripheral nerve schwannoma elicits an abscopal effect and synergizes with αPD1 treatment. **(A)** Schematic of bilateral *Nf2^-/-^* tumor implantation, systemic αPD1 treatment, and local radiation therapy to the right sciatic nerve schwannoma. **(B)** Tumor diameter measured by caliper. All animal studies are presented as mean±SEM, N=8 mice/group, and representative of at least three independent experiments. In vivo study significance was analyzed using a Student’s t-test. *P<0.01. ** P<0.001.

### RT of peripheral nerve schwannomas synergizes with αPD1 to control intracranial tumors and preserve hearing

Based on the abscopal effects observed, we next investigated whether local irradiation of the sciatic nerve tumor could synergize with systemic αPD1 treatment to inhibit intracranial tumor growth and prevent hearing loss. In this study, we implanted tumors in both the sciatic nerve and the CPA area. When the sciatic nerve tumors reached ∼5 mm in diameter, the mice were randomized into control or treatment groups of i) 5 Gy RT to the sciatic nerve tumor alone, ii) αPD1 alone, or iii) a combination of αPD1+5 Gy RT to the sciatic nerve tumor (Figure 5A). Mice were sacrificed when they exhibited ataxia, a symptom caused by the CPA tumors. Compared to the control group, local irradiation of the sciatic nerve tumor extended animal survival by 4 days (Figure 5B) and prevented tumor-induced hearing loss (Figure 5C). The RT abscopal effect synergized with systemic αPD1 treatment and significantly prolonged mice survival, with 66.7% of animals surviving over 100 days (Figure 5B). More importantly, we found that irradiation of the sciatic nerve tumor synergized with systemic αPD1 treatment and more effectively prevented tumor-induced hearing loss compared to αPD1 monotherapy (Figure 5D).

**Figure 5.**
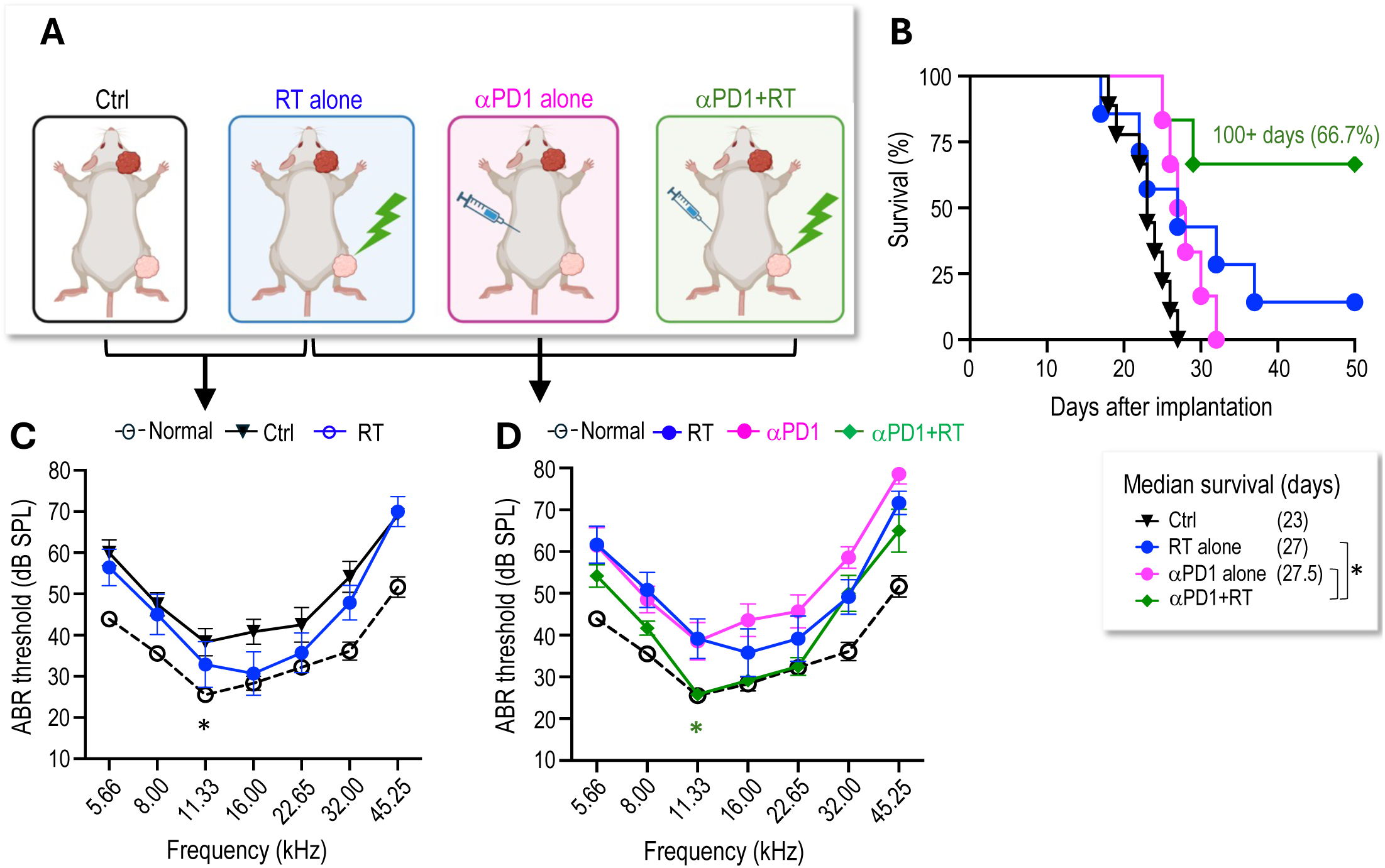
RT of peripheral nerve schwannomas synergizes with αPD1 to control intracranial tumors and preserve hearing. **(A)** Schematic of local irradiation of the sciatic nerve *Nf2^-/-^* tumor and systemic αPD1 treatment in mice bearing *Nf2^-/-^* CPA tumors. **(B)** Kaplan-Meir survival curve of mice. **(C)** ABR threshold of the ear ipsilateral to tumor implantation in non-tumor bearing (black dotted line), control (black line), and 5 Gy RT alone (blue line) groups. **(D)** ABR threshold of the ear ipsilateral to tumor implantation in non-tumor bearing mice (black dotted line), and in mice receiving 5Gy RT alone to the SN tumor (blue line), *i.p.* injection of αPD1 alone (pink line), or 5 Gy to the SN tumor + *i.p.* injection of αPD1 (green line). *P<0.01 for green vs. blue and green vs. pink. All animal studies are presented as mean±SEM, N=6 mice/group, and representative of at least three independent experiments. In vivo study significance was analyzed using a Student’s t-test. *P<0.01.

### Combined radiotherapy and αPD1 treatment activate dendritic cells and CD8 T cells

Compared to the αPD1 monotherapy, adding RT resulted in a significant increase of activated dendritic cells (DCs) and interferon (IFN)-γ-producing CD8^+^ T cells in the tumor, indicating enhanced antitumoral antigen presentation and activation of CD8 T cells (Figure 6A). The RT-induced increase in tumor-infiltrating T cells was further confirmed by immunofluorescent analysis (Supplemental Figure 2A). In the CPA tumors collected from mice in the combination group, we observed more apoptotic tumor cells (TUNEL^+^) and fewer proliferating tumor cells (PCNA^+^) compared to those in the control or monotherapy groups (Supplemental Figure 2B-C).

**Figure 6.**
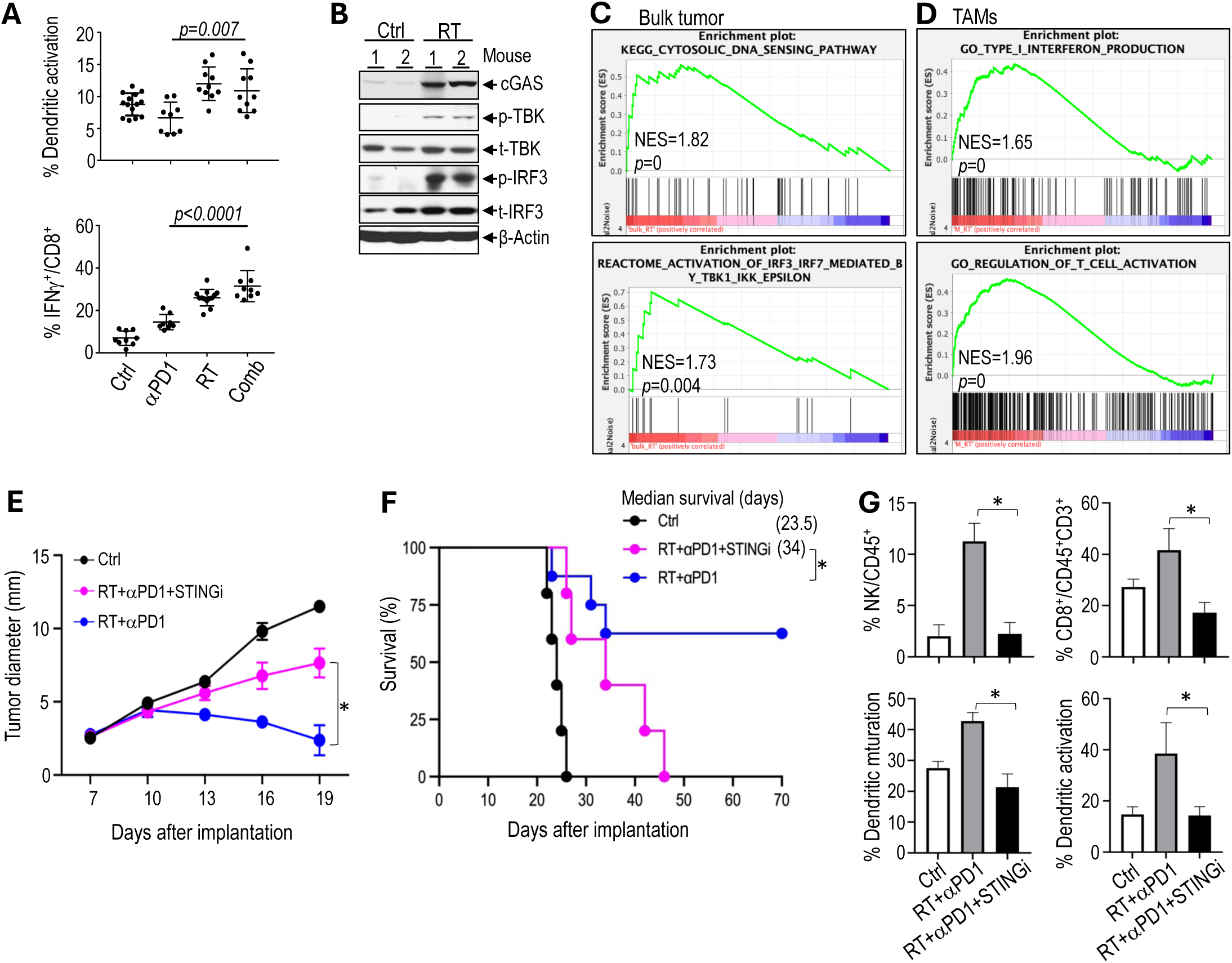
STING signaling mediates the synergistic effect observed in the combined RT and αPD1 treatment in Schwannoma models. **(A)** Flow cytometry analysis of activated dendritic cells and CD8^+^ T cells in *Nf2^-/-^* tumors. **(B)** Western blot of tumor tissues (N=2 tumors/group). **(C)** GSEA enrichment plots comparing irradiated vs. non-irradiated control bulk *Nf2^-/-^* tumors. **(D)** GSEA enrichment plots of TAMs isolated from irradiated vs. non-irradiated control *Nf2^-/-^* tumors. **(E)** Tumor diameter measured by caliper. **(F)** Kaplan-Meir survival curve of mice. **(G)** Flow cytometry analysis of NK cells, CD8 T cells, and dendritic cell maturation and activation in *Nf2^-/-^* tumors. All animal studies are presented as mean±SEM, N=8 mice/group, and representative of at least three independent experiments. In vivo study significance was analyzed using a Student’s t-test. Gene expression and flow cytometry data are presented as mean ± SD, and analyzed using Student’s t-test and the Mann-Whitney test. *P<0.01.

### STING signaling mediates the synergistic effect observed in the combined RT and αPD1 treatment in Schwannoma models

We next explore the molecular mechanisms that mediate the therapeutic benefit from the combination treatments. In the CPA tumors, we found that RT induced the expression of cyclic GMP-AMP (cGAMP) synthase (cGAS), phosphorylation of TANK-binding kinase 1 (TBK1) and interferon regulatory factor 3 (IRF3)(Figure 6B). RT-induced DNA damage activate the cytosolic DNA sensing pathway mediated by cGAS and stimulator of interferon genes (STING). STING recruits TBK1 and activates IRF3 to induce the expression of type I interferon (IFN), which activates the immune response ^[46, 47]^, and is critical for immune checkpoint inhibitor therapy and radiotherapy ^[48–51]^. Using RNASeq, we confirmed that RT induced the expression of genes i) in the cytosolic DNA sensing and IRF3-TBK1 pathways in the *Nf2^-/-^* bulk tumor (Figure 6C); and ii) in the type I IFN and T cell activation pathways in tumor-associated macrophages (TAMs) extracted from *Nf2^-/-^*tumors (Figure 6D). As the bulk tumor and single-cell RNASeq data converge on DNA sensing, IRF3-TBK, and IFN production, we focused on the STING pathway.

To investigate whether STING signaling is essential in mediating the therapeutic benefit of the combination treatment, we treated mice bearing *Nf2^-/-^*tumors in the sciatic nerve with i) control, ii) RT+αPD1, and iii) RT+αPD1+STING inhibitor (H151). We found that the addition of the STING inhibitor abolished the therapeutic benefits of the combination treatment: i) H151 significantly reduced the tumor growth delay achieved by the combination therapy (Figure 6E), ii) shortened median survival to 34 days, compared to the 62.5% long-term survival rate observed in the combination treatment group (Figure 6F), and iii) abolished the combination treatment-induced recruitment of immune effector NK cells and CD8^+^ T cells as well as the activation of dendritic cells (Figure 6G). These data suggests that STING signaling is essential for the therapeutic benefit of the RT and αPD1 combination treatment.

### Discussion and conclusion

In our studies, we demonstrated that the combination of αPD1 treatment with RT provides three significant benefits: i) Enhanced αPD1 efficacy and immune memory: RT induces immunogenic cell death and activates the STING pathway, enhancing αPD1 efficacy and generating long-term immune memory, ii) Reduced RT dose and associated tissue injury: The combination strategy reduces the required RT dose necessary for effective tumor control, potentially minimizing RT injury to surrounding normal tissues, and iii) Elicited abscopal effects on cerebellopontine angle (CPA) schwannomas: RT to peripheral nerve tumor induces a systemic abscopal effect, which synergizes with αPD-1 to effectively control intracranial schwannomas without direct irradiation, sparing the cochlea from radiation exposure and avoiding auditory radiation injury.

Our first finding is that RT, by inducing immunogenic cell death and activating the STING pathway, enhances the efficacy of αPD1 treatment. While immunotherapy has revolutionized cancer treatment, its efficacy as monotherapy remains limited, with ICIs benefiting fewer than 20-30% of patients with non-small cell lung cancer (NSCLC), renal cell carcinoma, or melanoma ^[52]^. To date, indications for ICIs have often combined treatment strategies to enhance efficacy. In addition to its direct tumoricidal effects, focal RT can trigger systemic antitumor immunity by: i) promoting *in situ* vaccination through the release of tumor-associated neoantigens ^[53]^, ii) activating dendritic cells to enhance cancer cell recognition by cytotoxic T lymphocytes (CTLs) ^[51]^, and iii) inducing immunogenic cell death ^[54]^. Combined RT and immunotherapy have shown increased efficacy in mouse models of colorectal cancer ^[55]^, breast cancer ^[56]^, pancreatic tumor ^[57]^; however, this strategy has not been evaluated in non-malignant schwannoma models.

In our VS model, we observed that RT induced immunogenic cell death, a form of regulated cell death that is sufficient to activate an adaptive immune response ^[58]^. Specifically, irradiation led to the release of immunostimulatory damage-associated molecular patterns (DAMPs), including ATP and HMGB1, from irradiated, dying tumor cells. At the molecular level, we found that RT activated the cGAS/STING pathway, leading to increased production of type I interferon, TNFα, and IL-1β. These RT-induced immune mediators support αPD1 therapy to enhance its efficacy. Despite the encouraging preclinical data on the improved efficacy of RT and αPD1 combination therapy, clinical studies have yielded mixed results. Maintenance αPD1 treatment following standard-of-care chemoradiotherapy has been shown to enhance overall survival (OS) in patients with NSCLC ^[59, 60]^; neoadjuvant stereotactic body radiotherapy (SBRT) combined with durvalumab (an αPD-L1 antibody) increases durvalumab response in early-stage NSCLC ^[61]^, while nivolumab (an αPD1 antibody) improves disease-free survival in NSCLC patients who received neoadjuvant chemoradiotherapy ^[62]^. Conversely, clinical studies in glioblastoma and head & neck squamous cell carcinomas failed to demonstrate a therapeutic benefit over RT or immunotherapy alone ^[63–66]^. Our findings provide a rationale for future clinical trials of this combination strategy in patients with VS, suggesting that optimally combining RT with immunotherapy may be a key for unlocking the potential of both therapies.

Secondly, we found that combined αPD1 treatment can help reduce the RT dose required for effective tumor control. RT is the standard of care for progressive or symptomatic vestibular schwannomas ^[67]^. Although fractionated radiation therapy approaches 100% tumor control, patients typically experience hearing deterioration within six months of treatment, suggesting that hearing loss results from RT-induced ototoxicity rather than tumor progression ^[41]^. Radiation therapy is known to cause permanent sensorineural hearing loss in a dose-dependent, progressive manner ^[68]^, and RT can also cause conductive hearing loss due to ear canal stenosis ^[69]^. Developing combination treatment regimens that reduce the RT dose while maintaining clinical efficacy is essential to minimize ototoxicity in VS therapies. In our VS model, we found that adding αPD1 treatment allowed for a reduction in the RT dose required to control tumor growth, suggesting that this combination strategy could help mitigate RT-induced ototoxicity. These findings support further clinical investigation of combined RT and αPD1 treatment in patients with VS to enhance therapeutic outcomes and preserve hearing function.

Lastly, as *NF2*-SWN patients often develop schwannomas throughout the body, we simultaneously implanted schwannoma cells in the CPA and in the sciatic nerve in mice. We observed that irradiation of a peripheral nerve tumor induced an abscopal effect, which synergized with systemic αPD1 treatment and effectively controlled intracranial tumor growth and, more importantly, prevented tumor-induced hearing loss without the need for direct irradiation of the CPA tumor. This strategy holds substantial clinical potential. Hypo-fractionated RT can elicit an abscopal effect, limiting the progression of distant tumors outside the locally irradiated area ^[70]^, and synergizing with ICI in both preclinical breast cancer, colorectal cancer, melanoma and lung cancer mouse models ^[56, 71–73]^, and lung cancer and melanoma patients ^[74, 75]^. The cochlea is particularly sensitive to RT compared to either the brain or the auditory nerve ^[76, 77]^. Direct irradiation of the cochlea can damage critical auditory structures, including the organ of Corti, basilar membrane, spiral ligament, and stria vascularis, as well as causing outer hair cell loss in the basal turn. This damage is most manisfested in the high-frequency hearing loss observed in patients ^[78]^. Our findings suggest that targeting peripheral nerve tumors while avoiding direct cochlear irradiation could reduce treatment-related side effects and better preserve hearing. This combination approach represents a promising shift toward more personalized, less toxic VS treatments. Future clinical trials in VS patients should optimize dosing and scheduling to maximize the therapy’s effectiveness. Additionally, extending this RT+αPD1 strategy to other solid tumors may help protect radiosensitive organs, thereby improving patient quality of life by minimizing treatment-related toxicities.

In summary, our studies demonstrate that combining αPD1 treatment with RT significantly improves the efficacy of both therapies. Combined RT enhances the efficacy of αPD1 therapy by promoting immunogenic cell death and activating STING pathway, while combined αPD1 therapy allows for a reduced RT dose, potentially minimizing RT-induced ototoxicity and other adverse effects. Most importantly, RT induces an abscopal effect that effectively controls intracranial tumors without directly irradiating the cochlea. This approach has the potential to protect the cochlea from radiation exposure, thereby preventing auditory damage and preserving hearing function. Our findings highlight the therapeutic potential of combining RT and αPD1 as a strategy to optimize tumor control while minimizing treatment-associated side effects, paving the way for more targeted and patient-centered cancer therapies.

## Notes

### Data availability

All data and the supplementary materials from this study are included in this manuscript and are available after publication upon request from the corresponding author. Sequencing data will be deposited in GEO and will be available upon publication.

## Acknowledgments

We thank Mark Duquette and Anna Khachatryan for their superb technical support, and Dr. Peigen Huang for assisting in animal studies.

## Author contributions

L.X. designed the research and supervised the research; Z.Y., S. L., L.W., D.W.B., J.C. performed mouse model studies; Z.Y., Y.S., B.X., A.P.J., performed flow cytometry and histology analysis; W.H. analyzed RNASeq data; Z.Y., B.X., performed patient sample histological analysis; L.X., Z.Y., S.L., L.W., A.M., analyzed data; L.X., S.R.P., K.S., H.A.S., and wrote the paper.

## Supplementary Figures

**Figure S1.**
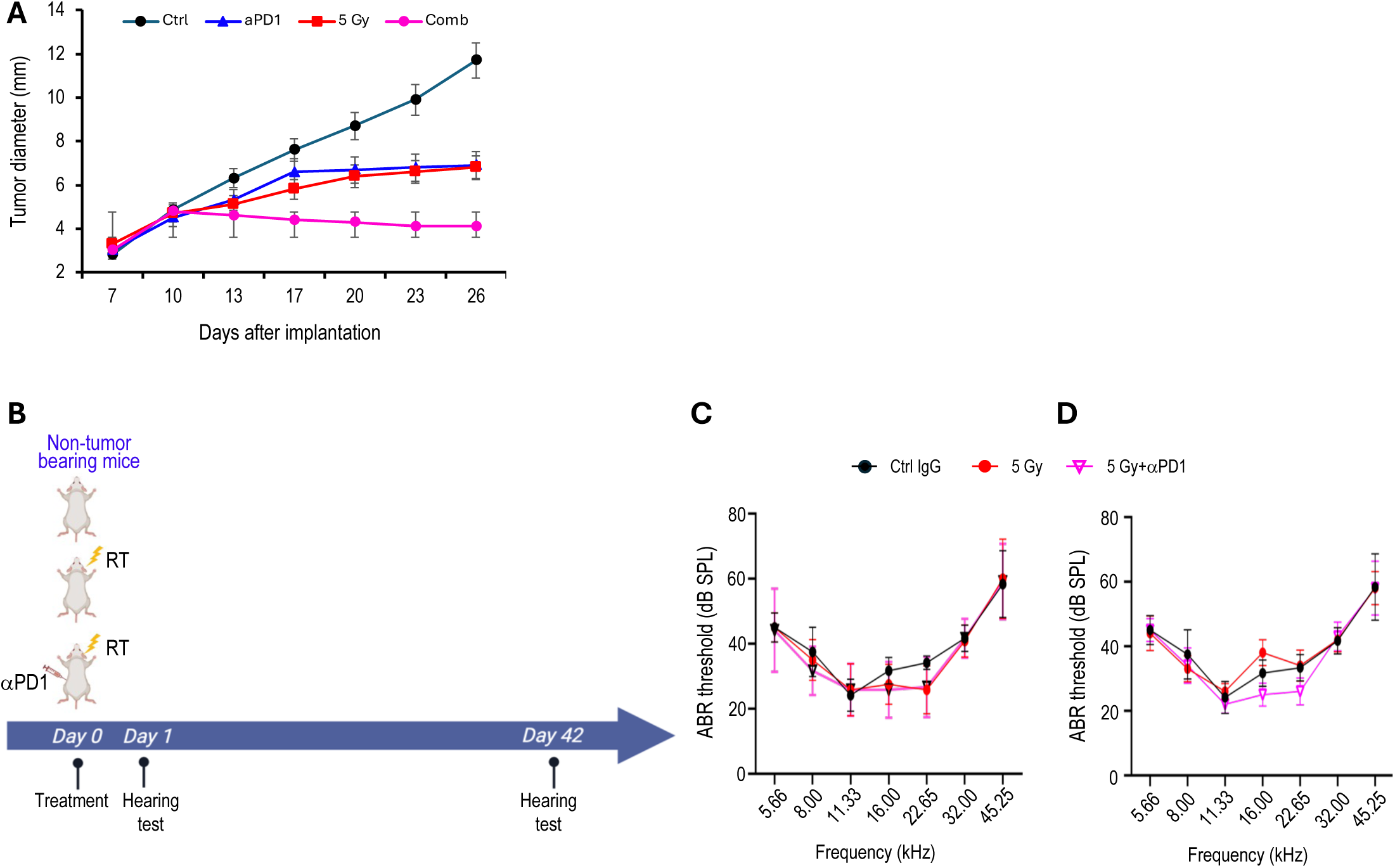
Combined αVEGF treatment enhances αPD1 efficacy in the mouse schwannoma models. **(A)** Tumor diameter measured by caliper in the SC4 SN schwannoma model. **(B)** Schematic and timeline of combined radiation therapy and aPD1 treatment in the CPA schwannoma models. **(C)** Day 1 post-treatment ABR threshold of mice bearing *Nf2^-/-^* tumor in the CPA model. **(D)** Day 42 post-treatment ABR threshold of mice bearing *Nf2^-/-^* tumor in the CPA model. All animal studies are presented as mean±SEM, N=6 mice/group, and representative of at least three independent experiments. In vivo study significance was analyzed using a Student’s t-test. *P<0.01, **P<0.001.

**Figure S2.**
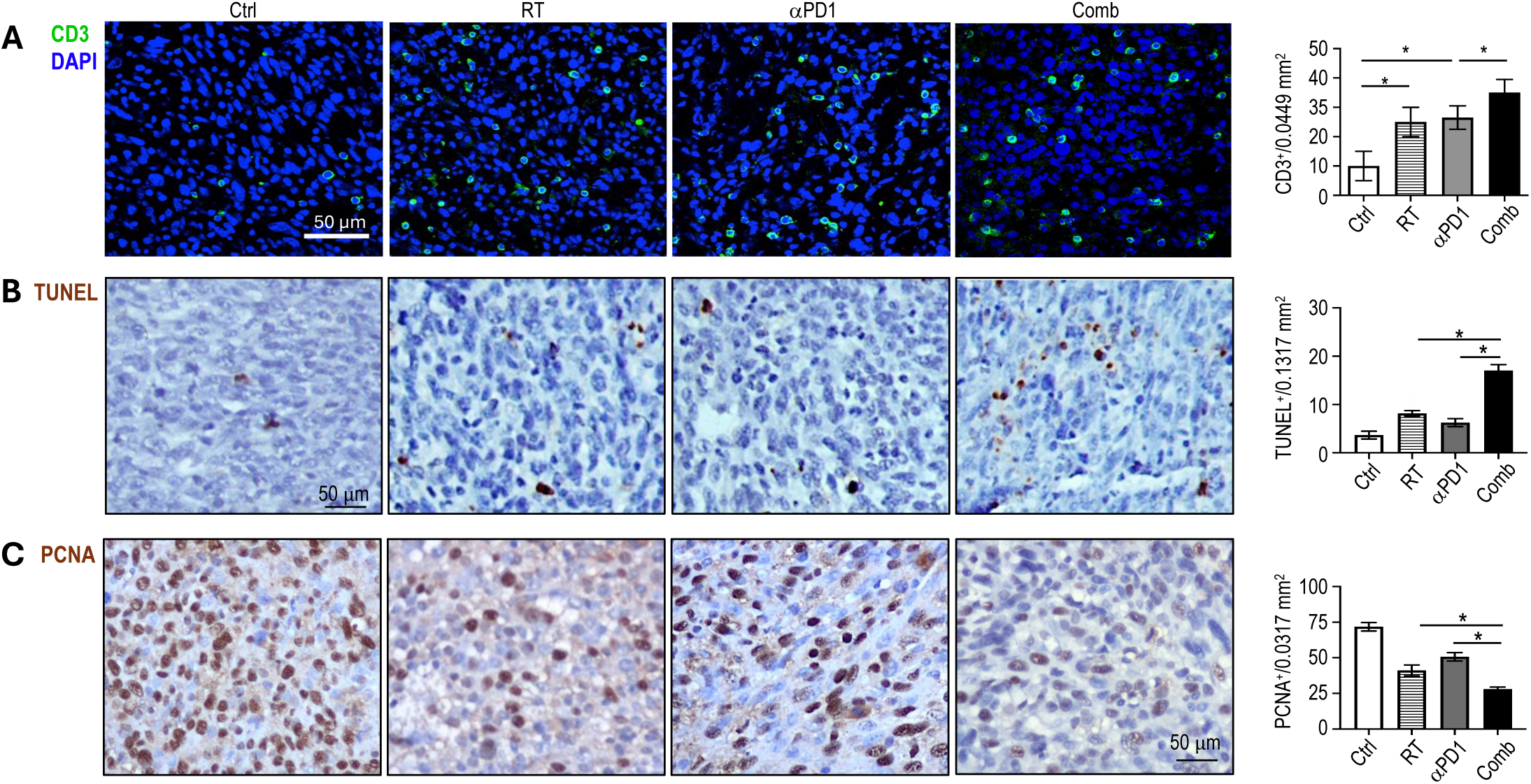
Combined radiotherapy and αPD1 treatment achieve enhanced efficacy in *Nf2^-/-^* model. **(A)** Representative immunofluorescent staining images of CD3 in *Nf2^-/-^* tumors. **(B)** Representative images of TUNEL staining for apoptotic cells **(C)** Representative images of PCNA staining for proliferating tumor cells. Image quantification and flow cytometry data are presented as mean ± SD, and analyzed using Student’s t-test and the Mann-Whitney test.

**Table S1.** Flow cytometry antibody panels.

| Surface Markers | Clone | Fluorophore | Catalog Number |
| --- | --- | --- | --- |
| CD11b | M1/70 | APC-Cy7 | BioLegend 101212 |
| CD11c | N418 | PE/Cy7 | BioLegend 117324 |
| CD206 | C068C2 | PE | BioLegend 141706 |
| CD3 | 17A2 | PE-cy7 | BioLegend 100219 |
| CD4 | GK1.5 | FITC | BioLegend 100405 |
| CD45 | 30-F11 | PerCP | BioLegend 103130 |
| CD8 | 53-6.7 | APC-cy7 | BioLegend 100713 |
| CD86 | GL1 | PE | BioLegend 105008 |
| F4/80 | BM8 | PE-Cy7 | BioLegend 123114 |
| Foxp3 | MF-14 | PE | BioLegend 126403 |
| Gr1 | RB6-8C5 | PE | BioLegend 108408 |
| Granzyme B | QA16A02 | APC | BioLegend 372204 |
| IFN-γ | XMG1.2 | PE/Dazzle594 | BioLegend 505846 |
| MHCII | M5/114.15.2 | APC | BioLegend 107614 |
| NK1.1 | S17016D | APC | BioLegend 156505 |
| TNF-α | MP6-XT22 | PE | BioLegend 506306 |

